# DeepSS2GO: protein function prediction from secondary structure

**DOI:** 10.1101/2024.03.30.584129

**Authors:** Fu V. Song, Jiaqi Su, Sixing Huang, Neng Zhang, Kaiyue Li, Ming Ni, Maofu Liao

## Abstract

Predicting protein function is crucial for understanding biological life processes, preventing diseases, and developing new drug targets. In recent years, methods based on sequence, structure, and biological networks for protein function annotation have been extensively researched. Although obtaining a protein in three-dimensional structure through experimental or computational methods enhances the accuracy of function prediction, the sheer volume of proteins sequenced by high-throughput technologies presents a significant challenge. To address this issue, we introduce a deep neural network model DeepSS2GO (Secondary Structure to Gene Ontology). It is a predictor incorporating secondary structure features along with primary sequence and homology information. The algorithm expertly combines the speed of sequence-based information with the accuracy of structure-based features, while streamlining the redundant data in primary sequences and bypassing the time-consuming challenges of tertiary structure analysis. The results show that the prediction performance surpasses state-ofthe-art algorithms. It has the ability to predict key functions by effectively utilizing secondary structure information, rather than broadly predicting general Gene Ontology terms. Additionally, DeepSS2GO predicts five times faster than advanced algorithms, making it highly applicable to massive sequencing data. The source code and trained models are available at https://github.com/orca233/DeepSS2GO.

## INTRODUCTION

Proteins are vital for a wide range of biological processes, serving key roles in cellular functions across both prokaryotic and eukaryotic organisms. An in-depth understanding of protein function not only has a considerable impact on meeting the academic demand for life science, but also drives advancements in the field of biomedicine(1). Protein function annotation can be achieved through biochemical experiments or computational methods. While the former is the gold standard due to its high accuracy and reliability, it is costly and low-throughput, making it unsuitable for the vast amount of protein sequence data generated by high-throughput sequencing instruments(2). Therefore, there is a pressing need to develop theoretical computational methods that combine accuracy with efficiency in protein function prediction(3).

Currently, there are multiple protein function classification standards, including Gene Ontology (GO)(4), EC(5), KEGG(6), and Pfam(7, 8), etc. Among these, the GO database is widely recognized for its relative comprehensiveness and systematic approach to describing the biological aspects of Molecular Function Ontology (MFO), Cellular Component Ontology (CCO), and Biological Process Ontology (BPO). The GO database is structured as a directed acyclic graph (DAG) with ‘is-a’ or ‘part-of’ relationships between GO terms, efficiently capturing protein functional characteristics(4). Meanwhile, if a protein is annotated with a GO term, all its ancestor terms up to the root of the ontology should also be annotated.

Protein function prediction methods can be categorized by the information utilized or the algorithms employed(9, 10). Information-based categorization includes methods grounded on primary sequence, tertiary structure, or proteinprotein interaction (PPI)(11). Algorithm-based categorization includes methods relying on sequence homology alignment (e.g. BLAST(12, 13), InterProScan(14), Multiple Sequence Alignment (MSA)(15), and Position-Specific Scoring Matrix (PSSM)(16)) and those based on deep learning (e.g. Convolutional Neural Networks (CNN)(17), Graph Neural Network (GNN)(18), Diffusion Network(19), Transformer(20, 21), and Large Language Models(22), etc). These two classification schemes intersect with each other. For instance, methods that rely on extracting primary sequence features can utilize various techniques, such as DeepPPISP(23) with PSSM, DeepGOPlus(24) employing CNN, and TALE(25) with transformer architecture. Additionally, methods like Graph2GO(26) and DeepFRI(27), which utilize three-dimensional (3D) structure leverage GNN, since the PDB is accessible. Furthermore, NetGO(28) harnesses PPI network information from the STRING database(29) for protein function prediction. Lastly, researchers often integrate multiple sources of information, such as combining protein sequence features with protein network information, as seen in the DeepGraphGO(30) approach.

Determining the relevant biological data and extracting essential features for model training is crucial beyond the variety of algorithms. This process is fundamental in leveraging biological information effectively. The essence of protein function prediction lies in learning the relationship between various biological information features and known functional labels within established species. This process is not about creating new GO terms; the predicted functions are inherently part of the existing functional pool. When confronted with unknown proteins, the trained model is employed in conjunction with the biological features of these proteins to score all the GO terms in the functional pool. The biological features of primary sequences encapsulate the patterns of the 20 amino acids. Different lengths and sequence orders correspond to diverse functions. Conversely, the biological features associated with tertiary structures pertain to spatial shapes, where distinct shapes and size features reveal critical insights and identify different functions.

According to the thermodynamic hypothesis proposed by Christian Anfinsen(31), the amino acid sequence dictates the protein tertiary structure, which is directly linked to its function. Therefore, incorporating 3D structural features is expected to improve the accuracy of protein function annotation, and many algorithms have also demonstrated this(26, 27). However, limitations of introducing 3D structures persist. While wet laboratory methods for analyzing protein 3D structures deliver accurate and reliable outcomes (32, 33), they are not sufficient to accommodate the demands of predicting functions from a large influx of new sequences(34), also entailing substantial financial and temporal costs (35). While computational methods like AlphaFold2(36) and trRosetta(37) have reduced this prediction time to a matter of hours, 3D-based function prediction methods still fall short of efficiently handling the vast amount of sequencing data produced by high-throughput sequencing instruments. Consequently, although methods that combine multiple sources of information typically outperform those solely relying on sequence data, obtaining these additional pieces of information in a short time frame is challenging. It may not be applicable to predict the function of less-studied proteins.

In cases where an abundance of primary sequences is available but tertiary structures are lacking, secondary structures exhibit distinct advantages. While primary sequences may vary significantly among different species, secondary structures, by eliminating redundant information, allow for a more focused investigation into the arrangement patterns of modules.

Secondary structures can be classified into eight categories according to the Dictionary of Secondary Structure of Proteins (DSSP)(38, 39): alpha-helix (H), 310-helix (G), pi-helix (I), beta-strand (E), beta-bridge (B), hydrogen-bonded turn (T), bend (S), and Coil (C). Although some research integrates protein secondary structure information as input features, like GAT-GO(40) and SSEalign(41), the application of secondary structure information has historically been constrained by data quality and quantity. We employ modified SPOT-1D-LM suite(42) to predict secondary structures for 68,325 primary sequences from SwissProt including all species in 20 hours, offering accuracy and speed suitable for processing large-scale datasets. Details will be illustrated in the Materials and Methods section in Table 1.

**Table 1.**
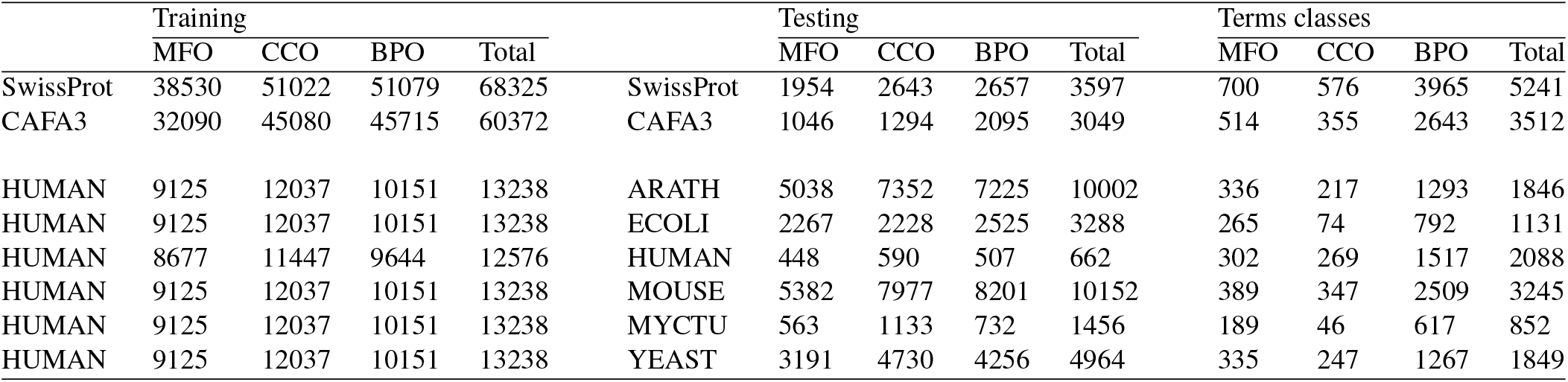
The number of protein sequences in the training-testing sets and the number of GO term classes grouped by sub-ontologies. Datasets include SwissProt, CAFA3, and training HUMAN testing other species. Detailed information on training one species and testing other species can be found in Supplementary Table S1.

Notably, relying solely on secondary structure predictions remains insufficient for protein function prediction, because of two reasons: First, this is due to variations in the amino acid arrangement of similar *α* helices (H, G, I) and *β* sheets (E, B), resulting in differences in physicochemical properties like charge and hydrophobicity. Second, intrinsically disordered protein regions, which play a critical role in alternative splicing, folding, and catalytic reactions(43), will be indiscriminately labeled with the secondary structure tag of Coil (C), thus failing to precisely differentiate between various functions. Consequently, we require the primary sequence and the homology information obtained through the Diamond algorithm(44) as a more refined supplement to the coarse secondary structure feature.

In this study, we propose a protein function predictor DeepSS2GO (Secondary Structure to Gene Ontology). This method utilizes deep learning models to extract features from the secondary structure and supplements them with primary sequence and homology information. This algorithm combines the sequence-based speed with the structure-based accuracy, while also overcoming shortcomings, like streamlining the redundant information in primary sequences and avoiding the time-consuming issue associated with tertiary structure analysis. When compared to similar algorithms, DeepSS2GO demonstrates superior performance in the MFO and CCO, while achieving second place in the BPO on both Maximum F-measure (*F*_max_) and Area Under the Precision-Recall Curve (AUPR) evaluation criterion. Furthermore, it attains the highest ranking in all three sub-ontologies when evaluated using the Minimum Sensitivity Index (*S*_min_) criterion. Notably, it accelerates computational speed by fivefold(45). Through two case analyses, we validate the efficacy of our approach in accurately predicting key functions of non-homologous proteins, providing comprehensive coverage. Moreover, the reduced training duration of our model facilitates prompt revisions of the SwissProt and GO databases, enhancing the timeliness of database updates.

## MATERIALS AND METHODS

### Overview

The overall concept and idea of the DeepSS2GO algorithm is illustrated in Figure 1. The primary sequence is analogous to fiber/gravel, the secondary structure to wooden/stone block, and the tertiary-quaternary structure to a wooden/stone bridge or tower. Traditional sequence-based prediction methods (model-aa) study the arrangement patterns of fiber and their relationship to the macroscopic object functional label as bridge or tower. However, due to potential differences in the arrangement of fibers and gravel, it might be difficult to predict whether it is a bridge or a tower. In situations where 3D spatial coordinates are not readily available, this study, instead of focusing on the arrangement of fibers, investigates the arrangement of intermediate wooden blocks and their relationship (model-ss8) to the macroscopic object functional label. Theoretically, compared to primary sequences, the model trained on secondary structures possesses greater translational capability, especially for crossspecies predictions.

**Fig. 1.**
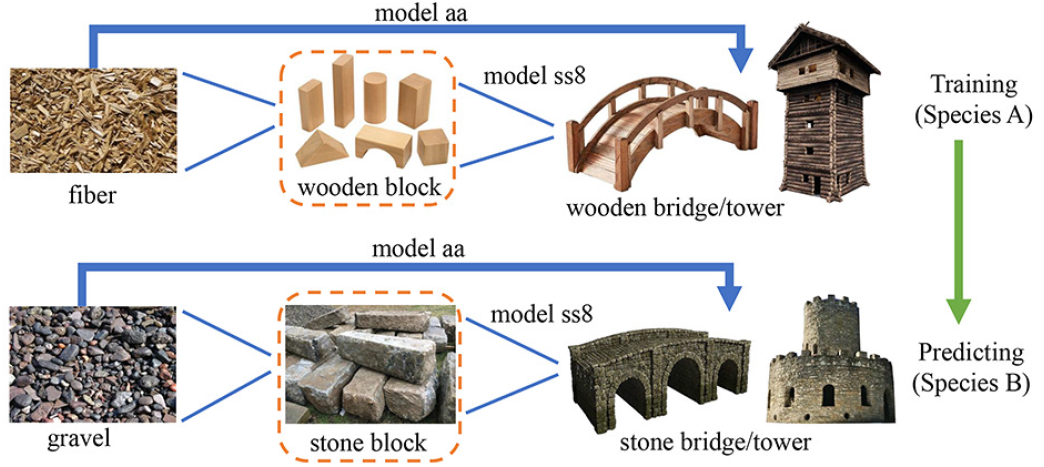
Overview of the DeepSS2GO conceptual diagram. This concept parallels two sets: ‘fiber wooden block wooden bridge’ and ‘gravel - stone block - stone bridge’, comparing them to the protein primary sequence - secondary structure - tertiary/quaternary structures. Traditional methods predict functions (bridge or tower), by studying the arrangement patterns of fiber or gravel (primary sequence features). This study introduces a new approach by examining the arrangement patterns of wooden blocks (secondary structure features) to predict functionality.

### Datasets

In this study, the functional annotations were derived from the GO(4) (June 2023), encompassing a comprehensive dataset distributed across three domains, including 47,497 terms: MFO (12,480), CCO (4,474), and BPO (30,543). Referencing the experiences from other studies(24, 45, 46), to enhance training efficiency and prediction accuracy, our training approach focused exclusively on GO terms with a sufficient number of training samples (i.e., the same GO label appearing in *≥* 50 sequences). Furthermore, annotations were propagated utilizing the relationships within the GO hierarchy(46).

Two datasets were employed in this research: SwissProt (47) (April 2023) and Critical Assessment of Function Annotation Challenge (CAFA3) (24). SwissProt, a subset of the UniProt database, is meticulously curated and manually annotated. Protein sequences and GO annotations used in this study are collected from SwissProt, retaining only experimental GO annotations with evidence codes IDA, IPI, EXP, IGI, IMP, IEP, IC, or TA. A total of six major species are selected for cross-validation training-testing: Arabidopsis thaliana (ARATH, 10,002), Escherichia coli (ECOLI, 3,288), Homo sapiens (HUMAN, 13,238), Mus musculus (MOUSE, 10,152), Mycobacterium tuberculosis (MYCTU, 1,456), and Saccharomyces cerevisiae (YEAST, 4,964).

The selection of these six species was primarily based on two reasons: First, the choice was to include species with a relatively large number of sequences, ensuring an ample amount of data for training and testing. The six selected species are all ranked within the top ten in terms of total quantity. Second, it was important to select species that include both closely related and significantly divergent organisms. This approach helps to reduce bias in horizontal comparisons, demonstrating the superiority of the ss8 model over the aa model across various cross-validation evaluations. Among the chosen species, there are closely related ones (HUMAN VS. MOUSE, ECOLI VS. MYCTU), as well as those that are significantly divergent, such as animal (HUMAN/MOUSE) VS. plant (ARATH), and two prokaryotic VS. four eukaryotic species.

Table 1 illustrates detailed statistics of the training and testing sets for the three domains in GO, including the number of proteins and GO labels. The datasets include SwissProt, CAFA3, and training HUMAN testing other species. Detailed information on training one species and testing other species can be found in Supplementary Table S1. Additionally, to facilitate comparisons with other cutting-edge protein function prediction methodologies (46), CAFA3(24) dataset is used for both training-testing sequences and functional annotations. Modified SPOT-1D-LM algorithm (42) is employed to predict secondary structures from primary amino acid sequences. Given that this algorithm utilizes the ESM-1b (21) and Prottrans (48) pre-trained models, it is constrained by protein length (less than 1024). After screening, a total of 68,325 protein sequences for Swissprot and 60,372 for CAFA3 were retained.

### The architecture of DeepSS2GO

The overall architecture is shown in Figure 2A. DeepSS2GO comprises three modules: two deep learning modules, one focused on secondary structures and the other on primary sequences, and a third module oriented towards homology alignment.

**Fig. 2.**
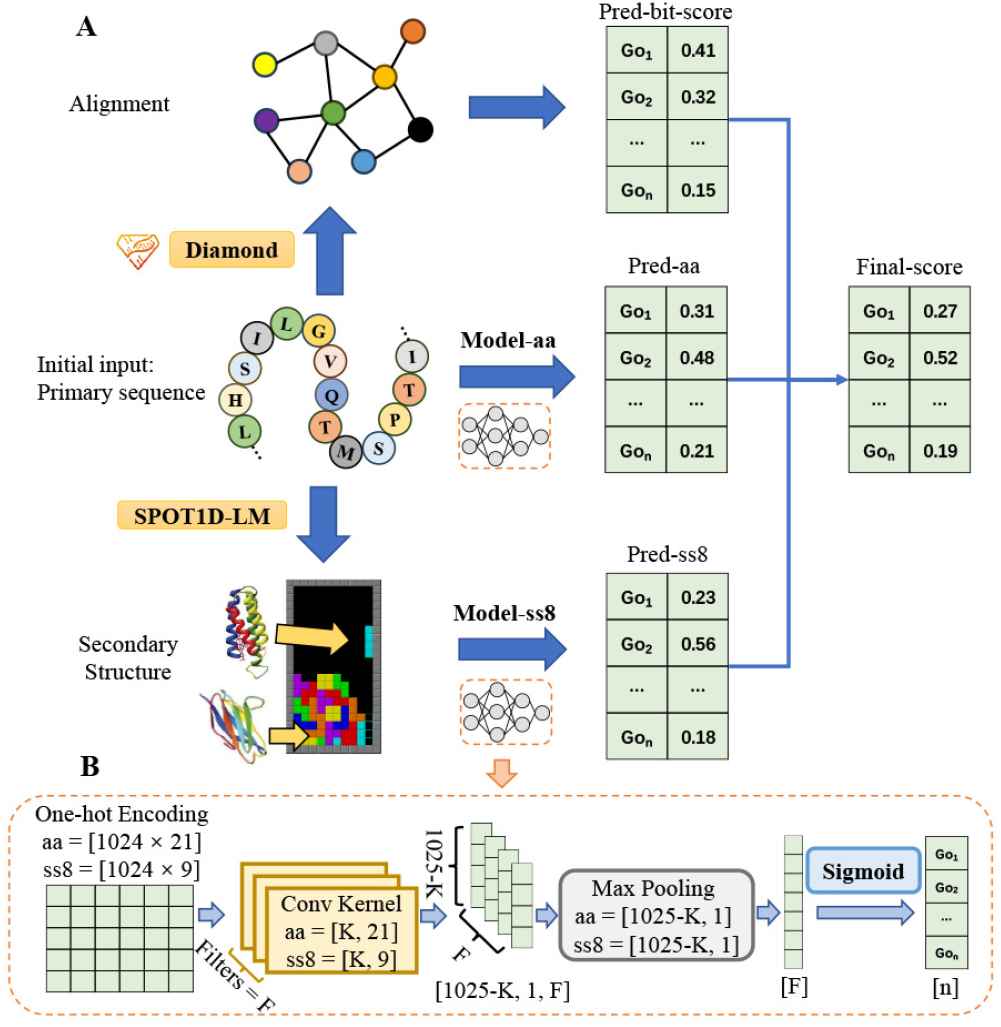
(A) The architecture of DeepSS2GO. The model consists of three components: a model trained on secondary structures (model-ss8), a model trained on primary sequences (model-aa), and Diamond homology alignment. First, the input primary sequence is converted into a secondary structure. Then, the primary sequence and secondary structure are separately processed through deep learning models to obtain preliminary predictions, Pred-aa and Pred-ss8. These, combined with the Pred-bit-score predicted by Diamond, are integrated to yield the Final-score. (B) The setup of model-aa and model-ss8. The input is a one-hot matrix, which passes through convolutional layers and pooling layers. After that, each term in the GO pool is scored individually using the Sigmoid activation function. For the one-hot matrices based on primary sequences and secondary structures, the sizes of the convolutional kernels and max pooling slightly differ. Kernel size ranges from 8 to 128 in increments of 8, while filter size ranges from 16 to 65536, doubling with each increment.

### Overall framework

The process begins by obtaining primary sequences and manually propagated annotations that have been filtered from the SwissProt training set, shown as the initial input in Figure 2A. Subsequently, data preprocessing is conducted. The altered SPOT-1D-LM suite is employed to convert primary amino acid sequences into secondary structures in bulk, *i*.*e*., replacing the original 20 amino acid letters with 8 letters representing secondary structures (H, G, I, E, B, T, S, C) (38, 39). Then, both the primary sequences and secondary structures are fed into the deep learning model (Figure 2B), respectively, yielding initial predictions for pred-aa and pred-ss8. On the other hand, homology comparison result Pred-bit-score is performed using the Diamond method(44), a remarkably high-speed and highperformance tool for conducting protein homology searches. The final prediction score is calculated by combining the three prediction scores (*S*_*aa*_, *S*_*ss*8_, and *S*_*Diamond*_) through Equation 1, where *α* and *β* are two hyperparameters balancing the influence of the three components, satisfying the conditions:0 *≤ α ≤* 1, 0 *≤ β ≤* 1, and 0 *≤ α* + *β ≤* 1.

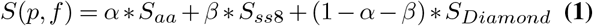

### Setup of Model

As both primary sequences and secondary structures are one-dimensional linear data structures, we employed the same deep-learning model for both. To highlight the advantages and effectiveness of secondary structures and to restore the biological essence as much as possible, we employ the most classic and concise convolutional neural network to extract their features.

We utilize PyTorch (49) to construct our neural network models, as depicted in Figure 2B. For a given protein sequence, we first convert the input primary sequence or secondary structure sequence into a one-hot matrix. If the input is a primary amino acid sequence, the matrix size will be [1024, 21], where the width 21 represents the 20 types of amino acids plus ‘other’. If the input is a secondary structure, the matrix size will be [1024, 9], where width 9 represents the 8 types of secondary structures plus ‘other’. 1024 is the length of the input, and sequences shorter than 1024 are padded with zeros. The input is then passed through a series of CNN layers with varying kernel sizes and filters, followed by Max Pooling layers, and normalized to the scoring range [0, 1] for n types of GO terms individually through the Sigmoid function. The training of a single model concludes within a maximum of 50 epochs. Additionally, we employed an EarlyStopping\ strategy with the patience of six epochs to prevent overfitting. Given the considerable parameter search space, after establishing certain hyperparameters such as the loss function (Binary Cross Entropy Loss), optimizer (Adam(50)), learning rate (0.0003), and activation function (Sigmoid), we focused on studying the parameters of kernel and filter size, which are more sensitive to protein function. These parameters will determine the features of specific sequences of particular sizes. Our model explores different combinations of kernels and filters, with kernel size varying between 8 and 128 in increments of 8, whereas the filter size ranges from 16 to 65536, doubling with each step.

The model-aa and model-ss8 are trained separately, and evaluated on the respective testing dataset for MFO, CCO, and BPO, thus determining the optimal kernel and filter size for each sub-ontology. The predicted GO scores by the best model, Pred-aa or Pred-ss8 will be combined with Predbit-score to yield the Final-score. Regarding the design of the model framework, we have also attempted to add fully connected layers after Max Pooling and represent secondary structures with three letters, but neither approach was satisfactory. Therefore, we will not elaborate further here.

### Implementation

We conduct two categories of experiments: specified cross-species testing, and testing that includes all species. To validate the enhanced translational ability of models trained on secondary structures, we employ cross-species testing, i.e., training with species A and testing with species B. To maximize primary sequence diversity, we aim to select species with significant distinctions, even using prokaryotic and eukaryotic organisms as separate training and testing sets. From SwissProt, six different species (ARATH, ECOLI, HUMAN, MOUSE, MYCTU, YEAST) are chosen for mutual testing. This selection includs two prokaryotes and four eukaryotes, the latter encompassing animals, plants, and fungi.

For the comprehensive species testing, the CAFA3 dataset is utilized for benchmarking against other similar algorithms, and the entire SwissProt dataset is employed to develop a model as complete and extensive as possible for predicting protein functions in new species. Specific details of the data can be found in section (Datasets).

If training and testing are conducted on the same species A (or the whole-species SwissProt dataset), then a random 5% of species A data is used as the testing set, with the remaining 95% as the training set. In contrast, if different species are used for training and testing, i.e., training on species A and testing on species B, then 100% of species A data is used as the training set, and 100% of species B data as the testing set. In both cases, 10% of the training set is allocated for validation during training. For instance, as shown in Table 1, there are a total of 13,238 HUMAN proteins and 10,002 ARATH proteins. If both training and testing are performed using HUMAN, then 12,576 proteins (95%) are used for the training set and 662 proteins (5%) for the testing set. If training HUMAN and testing ARATH, then all 13,238 HUMAN proteins are used as the training set, and all 10,002 ARATH proteins as the testing set. In total, we conducted 76 (i.e., 36 *×* 2 + 4) sets of tests, including cross-training/testing among the six species, and all-species dataset of SwissProt/CAFA3, for both aa and ss8. All training and testing processes are carried out on a Linux system equipped with a 24GB Nvidia GeForce RTX 3090.

### Evaluation metrics

Referring to relevant studies in this field(45, 46), we adapt three metrics for performance evaluation: *F*_max_, AUPR, and *S*_min_ (51–53). *F*_max_ is a metric that integrates precision and recall. The F-measure is the harmonic mean of precision and recall. *F*_max_ is the maximum F-measure achieved across all potential threshold settings, reflecting the optimal balance between precision and recall.

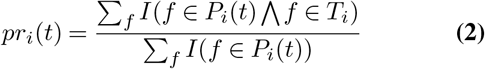

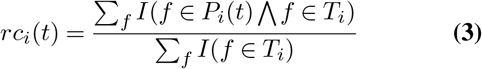

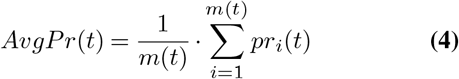

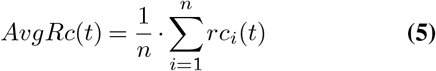

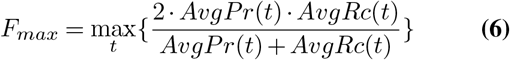

In this context, *f* represents a GO class, *Ti* denotes the set of true annotations, *P*_*i*_(*t*) refers to the predicted annotations set for a protein i at a specific threshold t, *m*(*t*) indicates the count of proteins that have at least one predicted category, *n* is the overall count of proteins, and *I* is a function that returns 1 if its given condition holds true, otherwise it returns 0. *AvgP r*(*t*) and *AvgRc*(*t*) represents the average precision and average recall at thresholds *t*, and calculated from *pr*_*i*_ and *rc*_*i*_ by the above formulas.

AUPR is the area under the precision-recall curve across all potential thresholds. It is a powerful tool for evaluating model performance in imbalanced datasets, especially when there is a substantial disparity in the number of positive and negative samples. Compared to the traditional Receiver Operating Characteristic Curve (ROC), AUPR is more sensitive to the predictive performance of a model for the minority class. This metric reflects a model’s ability to correctly identify positive (minority) instances amidst a large number of negatives (majority instances), focusing on precision and recall. In such contexts, AUPR is sensitive because it penalizes models more heavily for misclassifying the rare positive cases, thus providing a truer assessment of model performance on imbalanced datasets. It prioritizes the accurate detection of the minority class, highlighting the model’s effectiveness where it’s most needed.

*S*_min_, focusing on the minimum sensitivity index, a calculation of the gap between the true positive rate and the false positive rate across thresholds, sharply evaluates a classifier’s discriminative power between positive and negative instances. This metric is particularly insightful for assessing how well a model can differentiate between classes under varying conditions. A lower *S*_min_ indicates a model’s struggle to separate positive from negative cases effectively, often resulting in higher misclassification rates of crucial instances. In contrast, a higher *S*_min_ suggests that the model has a stronger capability to discern between the two, thereby reducing the likelihood of false positives and negatives. This sensitivity makes *S*_min_ an invaluable tool for model evaluation, especially in scenarios where the cost of misclassification is high. It pushes for models that not only recognize patterns but do so with a precision that minimizes the overlap between class distributions, enhancing the reliability of predictions in practical applications.

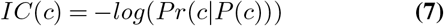

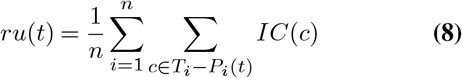

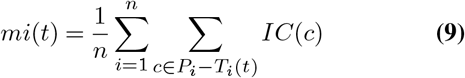

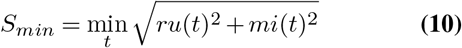

The information content, *IC*(*c*), is determined by the likelihood of annotations for class *c*. Here, *P* (*c*) represents the collection of parent classes for class *c*. Smin is derived using the equations below, where *ru*(*t*) represents the average residual uncertainty, and *mi*(*t*) denotes the average misinformation.

## RESULTS

This section encompasses the following aspects: Firstly, we validate the superiority of secondary structures over primary sequences in predicting functions by conducting cross-training predictions on proteins from different species. Secondly, we compare DeepSS2GO with other state-of-the-art methods, demonstrating the accuracy, efficiency, and updating convenience of our algorithm. Thirdly, we perform ablation experiments on the techniques used in DeepSS2GO.

Finally, we conduct two case studies to verify the effectiveness, efficiency, and comprehensiveness of the algorithm in predicting key functions.

### Superiority of Secondary Structures

Using the training and testing of all SwissProt data as an example, each training set employs either primary amino acid sequences or secondary structures as inputs. After training and evaluation, we can obtain results derived from primary amino acid sequences (see Supplementary Figure S1) and results stemming from secondary structures (see Supplementary Figure S2). Each figure comprises nine subfigures representing the evaluation results of three parameters: *F*_max_, AUPR, and *S*_min_, across three sub-ontologies: MFO, CCO, and BPO. The horizontal axis represents the logarithmic value of the filter size, and the vertical axis corresponds to the parameter values, with each plot representing the same kernel size. The extremum values of these three metrics can be found in the first two rows of Table 3.

For any fixed kernel, both *F*_max_ and AUPR values first increase and then decrease as the filter size rises. Reduction usually indicates overfitting. In the analysis with primary amino acid sequences, the peak *F*_max_ values observed for MFO, CCO, and BPO stand at 0.528, 0.666, and 0.426, respectively. These are achieved with kernel 16 and filter 32,768. When considering secondary structures, the maximum *F*_max_ values for MFO and BPO reach 0.616 and 0.452, respectively, both realized with kernel 32 and filter 32,768. In contrast, CCO highest *F*_max_ of 0.664 is attained with kernel 48 and filter 16,384. Comparatively, the model relying on secondary structures shows superior *F*_max_ values, exceeding the primary sequence model by 16.7% and 6.1% in MFO and BPO sub-ontologies, while matching performance in CCO.

Similarly, in AUPR, the secondary structure algorithm outperforms the primary sequence by 19.6% and 9.3% in MFO and BPO, respectively, and is on par in CCO. In *S*_min_, the secondary structure algorithm is higher by 13.4%, 1.1%, and 3.2% in MFO, CCO, and BPO, respectively. It is evident that the model based on secondary structures is markedly more effective in predicting the actual functions of proteins (MFO and BPO) compared to the primary amino acid sequence model. The two models perform comparably in determining protein components (CCO).

In addition to testing the whole SwissProt dataset, we select six species from SwissProt for cross-training and testing, as introduced in section (Implementation). Extracting the highest *F*_max_ values from the cross-validation of six different species, we obtained Figure 3 (AUPR and *S*_min_ results can be found in Supplementary Figure S3 and S4). In Figure 3, subfigures A, C, E represent predictions based on primary amino acid sequences (aa), while B, D, F represent predictions based on secondary structures (ss8). The same color in the upper and lower figures corresponds to the same GO subontology, with subfigures A and B representing MFO, C and D representing CCO, and E and F representing BPO. In each figure, a darker color indicates a higher *F*_max_ value.

**Fig. 3.**
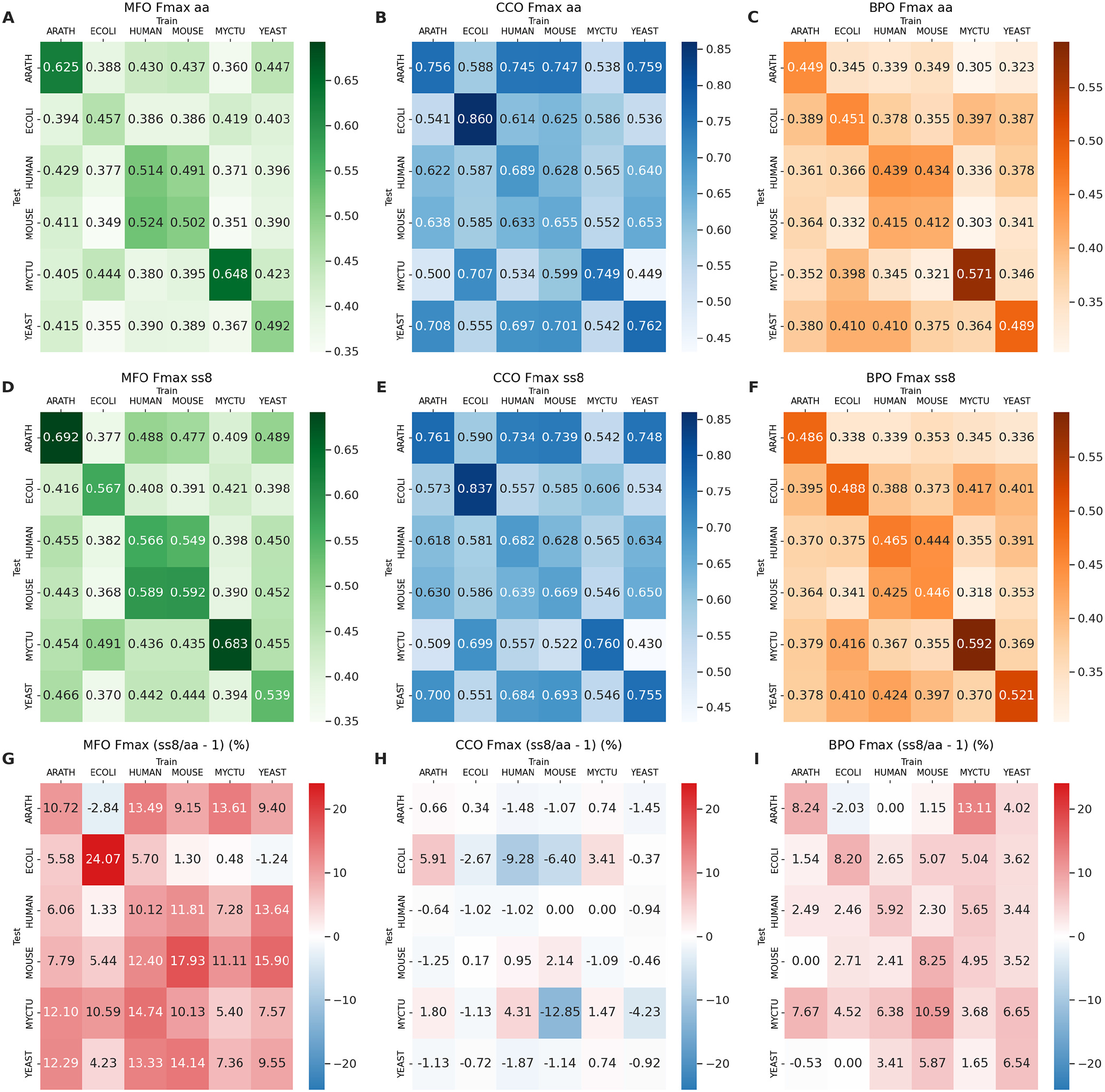
The performance of *F*_max_ scores in predicting GO functional annotations across species is depicted through heatmaps, utilizing models that have been trained and tested among six different species: Arabidopsis thaliana (ARATH), Escherichia coli (ECOLI), Homo sapiens (HUMAN), Mus musculus (MOUSE), Mycobacterium tuberculosis (MYCTU), and Saccharomyces cerevisiae (YEAST). (A) and (D) present MFO results based on model-aa and model-ss8 respectively; (B) and (E) show CCO results; and (C) and (F) illustrate BPO outcomes. Darker shades in the color gradients indicate higher metrics scores, reflecting greater prediction accuracy. Each matrix cell provides a metrics score for a model trained on the species denoted at the top and tested on the species labeled on the side. (G) depicts the percentage increased performance from model-aa to model-ss8 in MFO, similarly, (H) and (I) represent the increments in CCO and BPO, respectively. Red indicates the percentage of increase, while blue represents the percentage of decrease.

The following observations are noted: In examining the same subplot, aligning along the diagonal, the *F*_max_ values for selftesting consistently rank highest. Furthermore, the approach of training on eukaryotes and testing on prokaryotes demonstrates superior performance compared to the inverse. This disparity may be due to the more substantial sample size in eukaryotic training sets, potentially enhancing model accuracy.

Upon comparing the outcomes between aa and ss8, the subfigures G, H, and I of Figure 3 focus on the percentage increase in performance when transitioning from aa-trained models to ss-trained models within MFO, CCO, and BPO respectively. Red colors in the heatmaps signify a percentage increase in performance, while blue colors indicate a decrease. For MFO, ss8-*F*_max_ shows a notable improvement, approximately 5-20% higher than aa. This highlights the considerable advantage of secondary structures over primary sequences in the GO prediction of Molecular Function. Regarding CCO, ss8-*F*_max_ values are marginally lower, around 1-2% than those of aa. This indicates that aa encompasses more comprehensive CCO information density compared to ss8. In the context of BPO, ss8-*F*_max_ values generally outperform those of aa, with an increase of about 4-10%. An exception is observed in the scenario involving training on prokaryotes (ECOLI, YEAST) and testing on ARATH, where aa and ss8 yield comparable results.

The conclusions drawn from the AUPR and *S*_min_ (Supplementary Figures S3 and S4) analyses align with these observations. It is evident that secondary structures offer a clearer advantage in the prediction of protein functions. This is underlined by the fact that the structure dictates functions; secondary structures provide more structural information than primary sequences, enhancing their predictive capability for functions.

### Comparison with the state-of-the-art methods

Relying solely on primary sequences falls short in protein function prediction. To solve this issue, we integrate the equation 1 with secondary structure predictions, primary sequence data, and alignment score assessments for a holistic approach to function annotation. This section makes use of the CAFA3 dataset (24) for both training and testing components, facilitating comparative analysis of the DeepSS2GO algorithm against five other sequence-based methods: DiamondScore (24), DeepGOCNN (24), TALE+ (25), DeepGOPlus (24), and MMSMAPlus (46). The effectiveness of different algorithms is showcased through their best *F*_max_, AUPR, and *S*_min_ values across three sub-ontologies MFO, CCO, and BPO, as illustrated in Table 2.

**Table 2.**
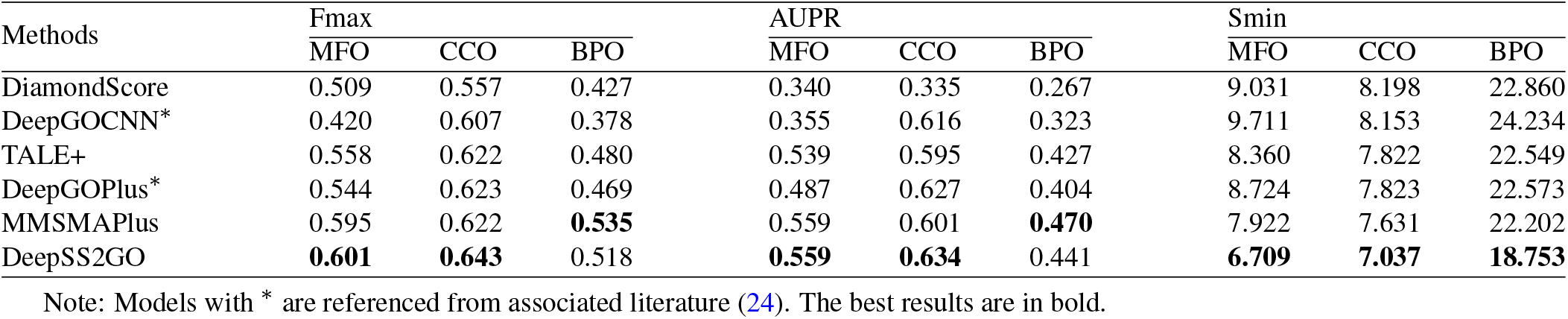
Performance comparison of DeepSS2GO against five state-of-the-art methods using the CAFA3 benchmark datasets.

| Methods | Fmax |  |  | AUPR |  |  | Smin |  |  |
| --- | --- | --- | --- | --- | --- | --- | --- | --- | --- |
|  | MFO | CCO | BPO | MFO | CCO | BPO | MFO | CCO | BPO |
| DiamondScore | 0.509 | 0.557 | 0.427 | 0.340 | 0.335 | 0.267 | 9.031 | 8.198 | 22.860 |
| DeepGOCNN* | 0.420 | 0.607 | 0.378 | 0.355 | 0.616 | 0.323 | 9.711 | 8.153 | 24.234 |
| TALE+ | 0.558 | 0.622 | 0.480 | 0.539 | 0.595 | 0.427 | 8.360 | 7.822 | 22.549 |
| DeepGOPlus* | 0.544 | 0.623 | 0.469 | 0.487 | 0.627 | 0.404 | 8.724 | 7.823 | 22.573 |
| MMSMAPlus | 0.595 | 0.622 | **0.535** | 0.559 | 0.601 | **0.470** | 7.922 | 7.631 | 22.202 |
| DeepSS2GO | **0.601** | **0.643** | 0.518 | **0.559** | **0.634** | 0.441 | **6.709** | **7.037** | **18.753** |

Note: Models with \* are referenced from associated literature (24). The best results are in bold.

Our model, despite leveraging the most traditional convolutional neural networks, still achieves excellent results, enhancing the protein function prediction performance on the CAFA3 dataset. Table 2 highlights DeepSS2GO superior performance in MFO and CCO for the metrics *F*_max_ and AUPR, and its near-best results in BPO. In terms of the *S*_min_ metric, it excels in all three sub-ontologies. The subsequent section (case 2), delves into the functional prediction of the protein LYPA2_MOUSE, emphasizing how the algorithm predicts specific sub-classes of GO functions and compares with other methodologies, underscoring the advantage of extracting features from protein secondary structures for predicting their functions.

Moreover, DeepSS2GO stands out for its high predictive accuracy coupled with computational efficiency. On average, the algorithm processes 1000 proteins from CAFA3 testing dataset in just 1.2 minutes, a substantive improvement over the cutting-edge algorithm, which requires approximately 7 minutes for 1000 proteins (45). This quintuple increase in speed is particularly beneficial when analyzing large volumes of sequenced, unknown metagenomic proteins.

In addition, the simplicity and user-friendliness of the model also mean that retraining costs are minimized. With continuous updates to the SwissProt and GO databases, our approach allows for rapid retraining to integrate new GO terms and discard outdated ones.

To sum up, the DeepSS2GO algorithm not only surpasses comparable methods in enhancing prediction performance on the CAFA3 dataset but also brings a substantial increase in processing speed. Furthermore, it provides a straightforward and update-friendly solution for adapting to the evolving landscape of protein and GO databases.

### Ablation study

We conduct ablation studies to demonstrate the efficacy of the three modules: aa, ss8, and Diamond, in the proposed DeepSS2GO framework (Figure 2A). Here, ‘aa’ symbolizes the model based on the primary amino acid sequence, and ‘ss8’ denotes the model based on the secondary structure. For a universally applicable validation model, we utilize the entire SwissProt database for both training and testing. Six sets of experiments were carried out, each involving different combinations of the three modules.

The findings deduced from the data presented in Table 3 can be summarized as follows: Initially, it is evident that the simultaneous presence of all three modules yields the best results, as highlighted in bold. The optimal values for *F*_max_ in MFO/CCO/BPO are 0.670, 0.703, and 0.535, respectively. For AUPR, the best values in MFO/CCO/BPO are 0.674, 0.742, and 0.493, respectively. Similarly, *S*_min_ achieves its optimal values in MFO/CCO/BPO at 7.682, 9.072, and 36.138, respectively.

**Table 3.**
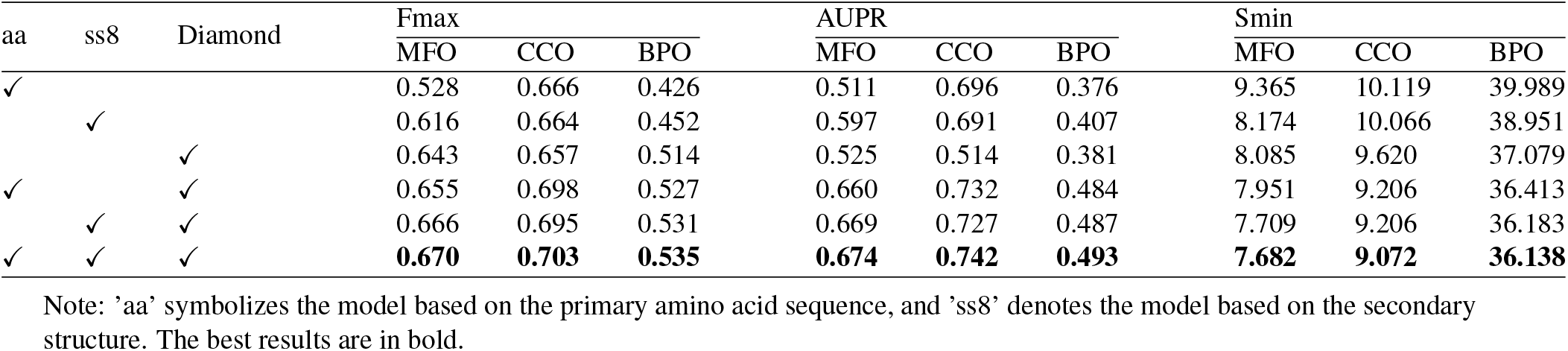
Assessment of the impact of different components within DeepSS2GO.

Furthermore, when only a single module is used, employing ss8 alone achieves the best AUPR scores, while Diamond alone performs best in terms of *F*_max_; the performance of aa alone is not as impressive. A comparison between the aa+Diamond and ss8+Diamond combinations reveals a slight edge for the latter.

Lastly, while using the Diamond alignment score alone shows some effectiveness, it is particularly valuable in complementing the deficiencies of either the model-aa or model-ss8, thereby enhancing the overall prediction accuracy. Thus, sequence homology information remains a precious source for functional inference.

### Case analysis

In this section, we conduct two sets of case studies. The first case is a self-comparison, where we demonstrate the superiority of features extracted from secondary structures over those from primary sequences by predicting the function of Surface Lipoprotein Assembly Modifier (SLAM) proteins. It shows that features from secondary structures better reflect the key functional GO terms and exhibit better generalizability in non-homologous proteins. The second case involves a horizontal comparison with similar methods, by predicting the function of the LYPA2_MOUSE protein. It proves that our predictor accurately predicts all the bottom-layer functionalities of the protein with high scores, and provides deeper and more precise GO annotations. Following communication with users of protein function prediction software, their usage habits have been understood. For novel, unknown proteins, researchers typically focus on the top 20-30 high-scoring results of GO terms in each of the three sub-ontologies as a preliminary judgment of the most probable functions. Therefore, the threshold does not have an absolute significance. Hence, in the following cases, the threshold is only used as a reference value for filtering.

### Case 1, Prediction of non-homologous SLAM proteins

SLAM1 and SLAM2 are two transport membrane proteins of the Neisseria meningitidis serogroup. Their primary function is to transport substrates, with SLAM1 targeting TbpB, fHbp, and LbpB, and SLAM2 targeting HpuA(54).

These two twin proteins are chosen as cases for several reasons. First, SLAM is not listed in the SwissProt database, thus not included in our training and testing sets. Second, homology comparisons of SLAM with other proteins in the SwissProt database yield a Diamond score of zero. This implies that SLAM is a non-homologous protein. Third, the sequence variance between the two proteins is substantial, with only about 25% sequence identity (Supplementary Figure S5B). However, their secondary structures are remarkably similar, each comprising one *β* barrel and multiple *α* helices (Supplementary Figure S5A), performing similar substrate transport functions. Therefore, these two SLAM cases effectively demonstrate that secondary-structure-based models are superior in predicting the function of non-homologous proteins with substantial sequence differences, compared to those primary-sequence-based models.

Three sets of tests are conducted using the aa+Diamond, ss8+Diamond, and aa+ss8+Diamond models to predict SLAM1 and SLAM2, with MFO results at a threshold of 0.06 presented in Table 4. The aa+Diamond model predicts broader, higher-level GO terms, but the inclusion of ss8 features allows for the prediction of specific terms such as GO:0005215, GO:0022857, GO:0022803, related to transporter and transmembrane activity. To be noticed, within the aa+Diamond module, SLAM1 exhibits scores of 0.011 for both GO:0005215 and GO:0022857, placing it at the 31st and 32nd positions in the list. Generally, researchers do not focus on GO terms that are ranked too low. In addition, in the case of SLAM2, also in the aa+Diamond module, there were no detected GO terms associated with transporter activity. Despite the low sequence similarity between SLAM1 and SLAM2, these GO terms are predicted due to the ss8model involvement, highlighting the transport-related functionalities essential to these proteins. Since SLAM proteins primarily act as substrate transporters, the highlighted parts in their functional annotations are particularly noteworthy.

**Table 4.**
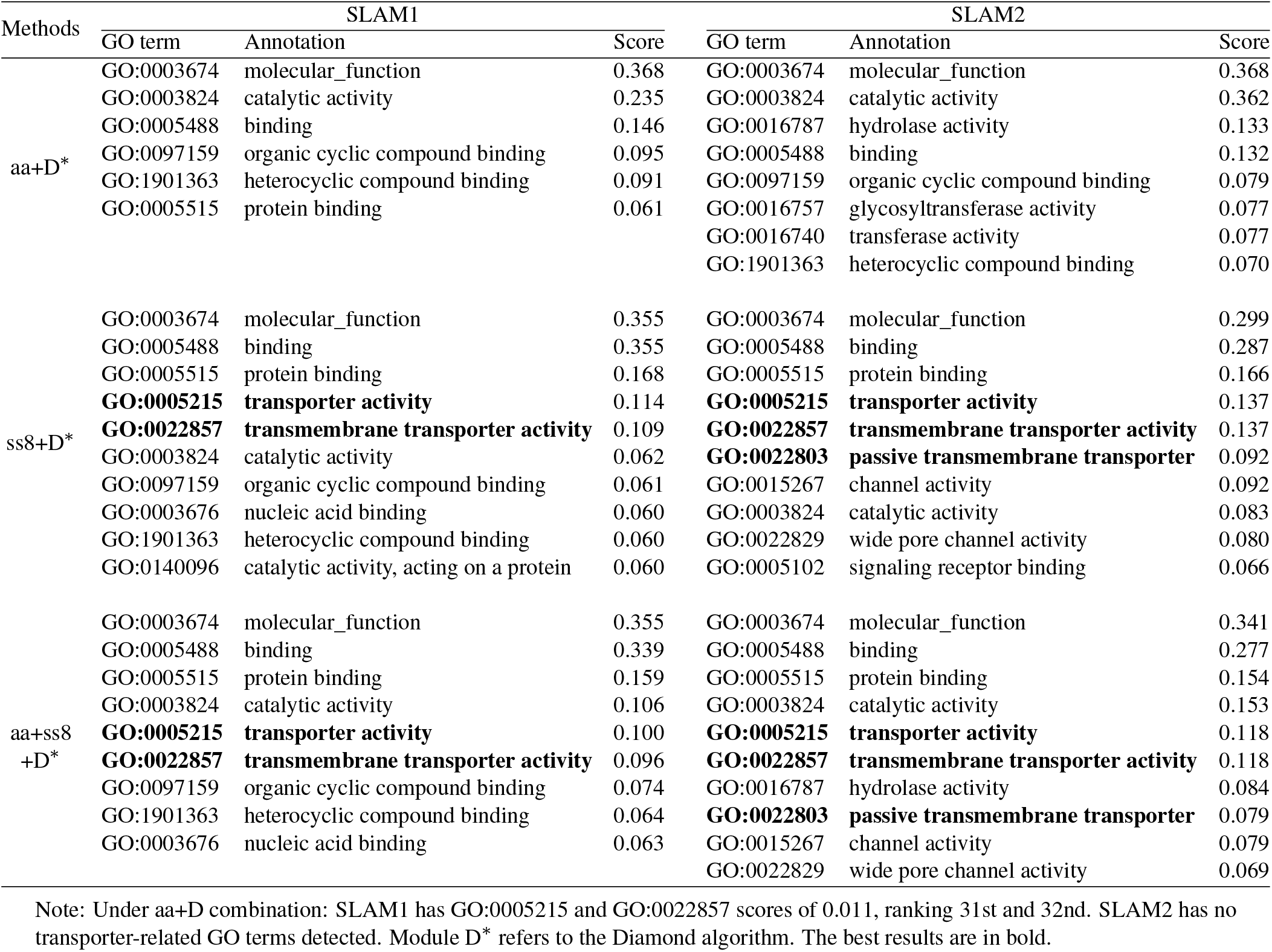
Evaluation of MFO prediction for SLAM1 and SLAM2 proteins, using different combinations of DeepSS2GO modules with a threshold of 0.06.

**Table 5.** Evaluation of the BPO prediction for the LYPA2_MOUSE protein, ranked in descending order of scores.

| GO term | Annotation | Score |
| --- | --- | --- |
| **GO:0008150** | biological_process | 0.870 |
| **GO:0008152** | metabolic process | 0.823 |
| **GO:0071704** | organic substance metabolic process | 0.814 |
| **GO:0009987** | cellular process | 0.781 |
| **GO:0044238** | primary metabolic process | 0.766 |
| **GO:0043170** | macromolecule metabolic process | 0.663 |
| **GO:0043412** | macromolecule modification | 0.663 |
| **GO:0098732** | macromolecule deacylation | 0.663 |
| **GO:0006807** | nitrogen compound metabolic | 0.654 |
| **GO:0019538** | protein metabolic process | 0.654 |
| **GO:0035601** | protein deacylation | 0.654 |
| **GO:0036211** | protein modification process | 0.654 |
| **GO:1901564** | organonitrogen compound metabolic | 0.654 |
| **GO:0009056** | catabolic process | 0.618 |
| **GO:1901575** | organic substance catabolic process | 0.614 |
| **GO:0042157** | lipoprotein metabolic process | 0.538 |
| **GO:0098734** | macromolecule depalmitoylation | 0.495 |
| **GO:0044237** | cellular metabolic process | 0.484 |
| **GO:1901565** | organonitrogen compound catabolic | 0.481 |
| **GO:0009057** | macromolecule catabolic process | 0.446 |
| GO:0050896* | response to stimulus | 0.444 |
| **GO:0030163** | protein catabolic process | 0.438 |
| GO:0006629* | lipid metabolic process | 0.413 |
| GO:0044255* | cellular lipid metabolic process | 0.413 |
| GO:0046486* | glycerolipid metabolic process | 0.413 |
| **GO:0002084** | protein depalmitoylation | 0.389 |
| **GO:0042159** | lipoprotein catabolic process | 0.389 |

Note: GO terms in bold are True Position predictions. Go terms with \* are referred to as False Positive predictions.

This case validates the hypothesis proposed in Figure 1. Despite the vastly different arrangement of fibers and gravels, the function of the macrostructure can be determined as a ‘bridge’ rather than a ‘tower’ by learning the arrangement patterns of blocks. Similarly, even with the diversity in primary sequences, the accuracy of protein function prediction can be improved by learning the arrangement patterns of secondary structures. Incorporating features extracted from secondary structures provides higher sensitivity in predicting protein functions compared to models based solely on primary amino acid sequences. Even though the three main evaluation metrics (*F*_max_, AUPR, *S*_min_) are indeed crucial for assessing the algorithm for general comparison, in a practical predicting application, it is vital to identify the specific key functions of an unknown protein. In this aspect, DeepSS2GO, which integrates secondary structure features, proves to be more effective.

### Case 2, Prediction of LYPA2_MOUSE protein

As the literature (46) has already examined the LYPA2_MOUSE protein (UniProt Symbol: Q9WTL7) and conducted comparisons with other similar algorithms, in this case, we also adapt this protein as our test object. The LYPA2_MOUSE protein serves as an acyl-protein thioesterase, responsible for hydrolyzing fatty acids attached to S-acylated cysteine residues in various proteins. A critical function of LYPA2_MOUSE includes facilitating the depalmitoylation process of zDHHC(55). Therefore, predicting depalmitoylation-associated GO terms (GO:0098734 and GO:0002084) is of crucial importance in understanding its biological process.

Since the LYPA2_MOUSE protein exists in the SwissProt training set, we remove this protein from the set, then retrain the model using the same kernel and filter parameters as the optimal solution, and predict this protein function with the new model. The BPO results are shown in Tabel 5. The DeepSS2GO algorithm successfully predicted all 23 GO terms with high scores, achieving an accuracy of 100%, highlighted in bold. Further analysis revealed that the success in predicting all GO terms mainly stemmed from accurately predicting the sub-node GO:0002084, which led to the inference of all parent-level label terms. Figure 4 compares the GO term labels predicted by DeepSS2GO with those by other similar algorithms. DeepSS2GO managed to accurately predict all labels, transcending all other same-type algorithms. This proves that our predictor provides deeper, more specific, and crucial functional annotations, making it a more practical method for the accurate and comprehensive prediction of protein functions in biological research.

**Fig. 4.**
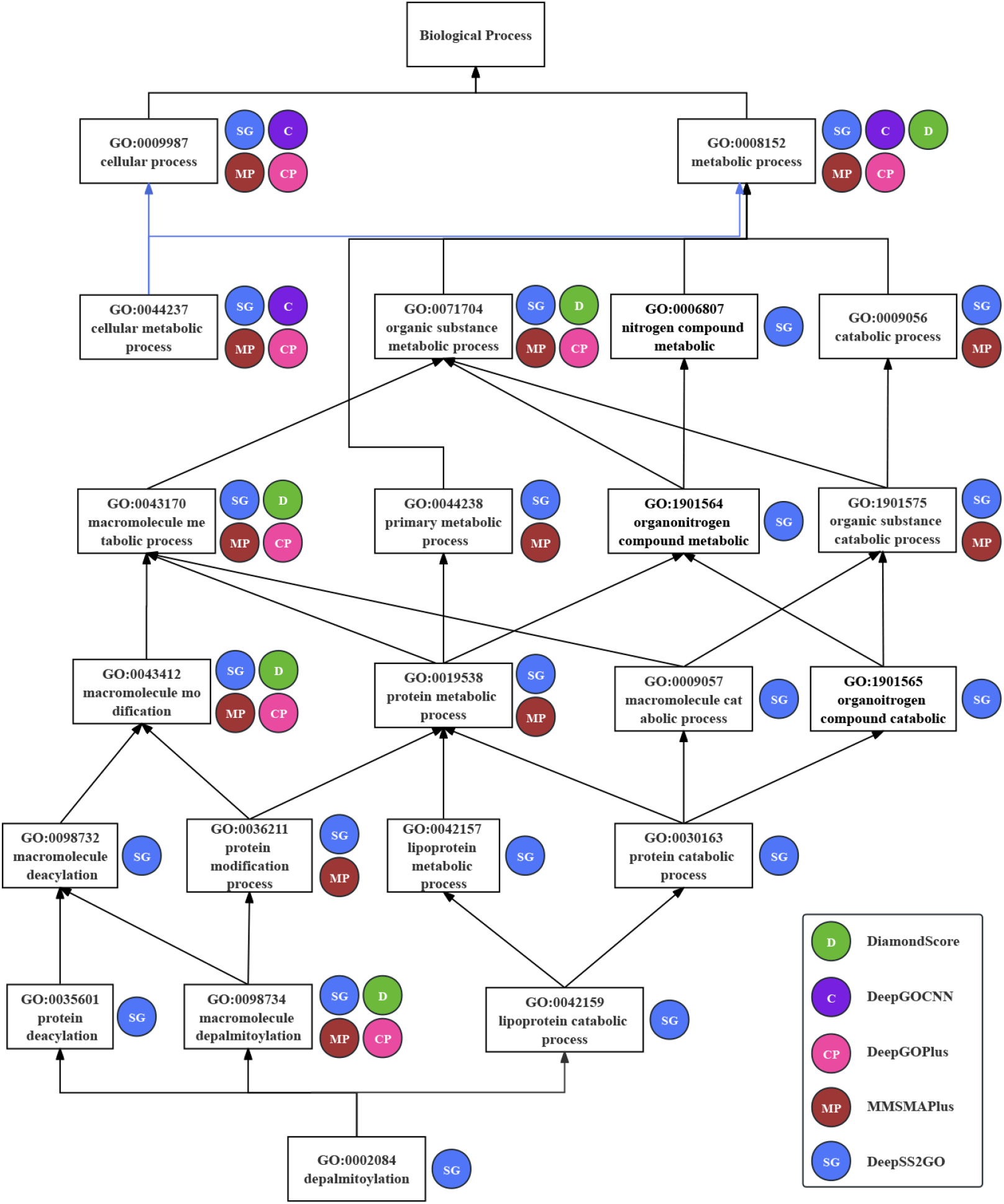
Comparison of predicted GO terms by various methods for LYPA2_MOUSE (UniProt Symbol: Q9WTL7) protein within the BPO directed acyclic graph. The established baseline for these predictions is derived from the propagation of experimental BPO annotations (GO:0002084).

It is important to note that some GO terms in the iterature(46) have been changed. For example, GO:0044260 is now obsolete, and GO:0044267 with GO:0006464 are secondary IDs for GO:0019538 and GO:0036211, respectively. Moreover, in the latest GO database used in this study, GO:0006807, GO:1901564, and GO:1901565 are defined as parent-level terms of GO:0002084(56). The following four aspects are continuously updated: the SwissProt database, the GO database, the interrelationships between GO terms, and the correspondence between proteins and GO terms. Therefore, the simplicity, time efficiency, and ease of retraining the DeepSS2GO model ensure regular updates.

These results indicate that the proposed DeepSS2GO method overtops the state-of-the-art methods in stability, reliability, accuracy, efficiency, and generalization to non-homologous proteins. It can further extend to proteins of species not ‘seen’ in the training set, predicting biological functions of new and unknown protein sequences.

## DISCUSSION

Protein function prediction methods based on primary sequences or tertiary structures exhibit inherent limitations. The information in primary sequences contains an overload of information, making it challenging to accurately predict functions from unknown species through amino acid sequence information alone. Although leveraging tertiary structure for function prediction improves accuracy, it is impractical for analyzing massive datasets due to its time-consuming nature. From primary to tertiary levels, it is precisely because the ‘functional information density’ continually increases that it becomes easier to predict function. This functional information density refers to the ratio of functional information to total information. Considering this, our developed secondary-structure-based prediction algorithm, DeepSS2GO, can compensate for these shortcomings, combining the efficiency of sequencing based on primary sequences with the accuracy of utilizing partial spatial structural information.

DeepSS2GO is characterized by its accuracy, critical insights, comprehensiveness, efficiency, and ease of updating. It enhances protein function prediction by reducing the redundant information of primary sequences through the modular integration of secondary structure features. This approach improves the prediction accuracy, relevancy, and breadth. Furthermore, DeepSS2GO outperforms current leading sequence-based predictors in performance, offering comprehensive predictions of essential protein functions and demonstrating excellent generalization capabilities for nonhomologous proteins and new species. Its rapid prediction capability makes it highly applicable in various fields, including metagenomics, for large-scale unknown species. Additionally, the user-friendly model architecture minimizes the costs associated with retraining, facilitating quicker and more convenient updates with the latest database.

However, there are areas where DeepSS2GO can be further improved. In an effort to emphasize the effectiveness of the secondary structure, the model was built using the classic conventional convolutional neural network, proving that even simple methods can yield outstanding results. Recent algorithmic developments in areas such as GNN(18), Diffusion mechanisms(45), Geometric Deep Learning(57), Selfsupervised learning(48), and Large Language Models(22) have shown exceptional utility in protein structure and function analysis. Applying these state-of-the-art algorithms to extract protein sequence information from various dimensions could enhance the accuracy of functional predictions. Moreover, the algorithm transition from primary to secondary sequence prediction uses ProtTrans and ESM pretrained models, which are limited by protein sequence length, thereby excluding large proteins over 1024 amino acids. Adopting more versatile secondary structure prediction methods for longer sequences in the future would expand our algorithm scope significantly. Lastly, functional prediction is not limited to full-length proteins but can also be applied to studying various polypeptides(58, 59), integrating multiple features which will facilitate a broader elucidation of disease mechanisms and the discovery of drug targets. Therefore, we aim to further integrate drug and disease information using information fusion methods, allowing functional annotation algorithms to be more effectively applied in practical applications, benefiting humanity.

Overall, DeepSS2GO combines advanced feature learning capabilities with cross-species transfer potential. As genomic sequencing progresses and the quantity of new species sequence data grows, this method promises to be a valuable tool for protein function prediction, balancing accuracy with computational efficiency.

## SUPPLEMENTARY DATA

Supplementary data are available online at https://academic.oup.com/bib

## DATA AVAILABILITY

The source code and trained models are available for research and non-commercial use at https://github.com/orca233/DeepSS2GO.

## Author contributions statement

F.S., M.N., and M.L. conceived and supervised the whole project. F.S. programmed the code and wrote the manuscript. J.S. performed data processing and analyzed the results. S.H. and N.Z. refined the algorithm and reviewed the manuscript. K.L. processed the case study. All the authors discussed, revised, and proofread the manuscript.

## Acknowledgments

The authors thank the HPC-Service Station, Cryo-EM Center, and Center for Computational Science and Engineering at Southern University of Science and Technology. M.L. is an investigator of SUSTech Institute for Biological Electron Microscopy.

