## supplementary_information for "DeepSS2GO: protein function prediction from secondary structure"

Fu V. Song[1,*], Jiaqi Su[1], Sixing Huang[2], Neng Zhang[3], Kaiyue Li[1], Ming Ni[4,*] and Maofu Liao[1,5,*]

[1] Department of Chemical Biology, School of Life Sciences, Southern University of Science and Technology, Xueyuan Avenue, 518055, Shenzhen, China
[2] Gemini Data Japan, Kitaku, Oujikamiya 1-11-11, 115-0043, Tokyo, Japan
[3] Electronic Engineering and Computer Science, Queen Mary University of London, Mile End Road, E1 4NS, London, UK
[4] MGI Tech, Beishan Industrial Zone, 518083, Shenzhen, China
[5] Institute for Biological Electron Microscopy, Southern University of Science and Technology, Xueyuan Avenue, 518055, Shenzhen, China

[*] Contact:

**Table S1.** The number of protein sequences in the training-testing sets and the number of term classes grouped by sub-ontologies. Datasets include SwissProt, CAFA3, and training one species testing other species.

| | Train | | | | | Test | | | | Terms classes | | | |
|---|---|---|---|---|---|---|---|---|---|---|---|---|---|
| | MFO | CCO | BPO | Total | | MFO | CCO | BPO | Total | MFO | CCO | BPO | Total |
| SwissProt | 38530 | 51022 | 51079 | 68325 | SwissProt | 1954 | 2643 | 2657 | 3597 | 700 | 576 | 3965 | 5241 |
| CAFA3 | 32090 | 45080 | 45715 | 60372 | CAFA3 | 1046 | 1294 | 2095 | 3049 | 514 | 355 | 2643 | 3512 |
| ARATH | 4790 | 6977 | 6879 | 9501 | ARATH | 248 | 375 | 346 | 501 | 121 | 114 | 643 | 878 |
| ARATH | 5038 | 7352 | 7225 | 10002 | ECOLI | 2267 | 2228 | 2525 | 3288 | 171 | 62 | 508 | 741 |
| ARATH | 5038 | 7352 | 7225 | 10002 | HUMAN | 9125 | 12037 | 10151 | 13238 | 336 | 217 | 1293 | 1846 |
| ARATH | 5038 | 7352 | 7225 | 10002 | MOUSE | 5382 | 7977 | 8201 | 10152 | 253 | 166 | 1096 | 1515 |
| ARATH | 5038 | 7352 | 7225 | 10002 | MYCTU | 563 | 1133 | 732 | 1456 | 119 | 38 | 407 | 564 |
| ARATH | 5038 | 7352 | 7225 | 10002 | YEAST | 3191 | 4730 | 4256 | 4964 | 189 | 167 | 803 | 1159 |
| ECOLI | 2267 | 2228 | 2525 | 3288 | ARATH | 5038 | 7352 | 7225 | 10002 | 171 | 62 | 508 | 741 |
| ECOLI | 2167 | 2130 | 2406 | 3123 | ECOLI | 100 | 98 | 119 | 165 | 89 | 34 | 233 | 356 |
| ECOLI | 2267 | 2228 | 2525 | 3288 | HUMAN | 9125 | 12037 | 10151 | 13238 | 265 | 74 | 792 | 1131 |
| ECOLI | 2267 | 2228 | 2525 | 3288 | MOUSE | 5382 | 7977 | 8201 | 10152 | 205 | 56 | 688 | 949 |
| ECOLI | 2267 | 2228 | 2525 | 3288 | MYCTU | 563 | 1133 | 732 | 1456 | 94 | 30 | 281 | 405 |
| ECOLI | 2267 | 2228 | 2525 | 3288 | YEAST | 3191 | 4730 | 4256 | 4964 | 160 | 59 | 507 | 726 |
| HUMAN | 9125 | 12037 | 10151 | 13238 | ARATH | 5038 | 7352 | 7225 | 10002 | 336 | 217 | 1293 | 1846 |
| HUMAN | 9125 | 12037 | 10151 | 13238 | ECOLI | 2267 | 2228 | 2525 | 3288 | 265 | 74 | 792 | 1131 |
| HUMAN | 8677 | 11447 | 9644 | 12576 | HUMAN | 448 | 590 | 507 | 662 | 302 | 269 | 1517 | 2088 |
| HUMAN | 9125 | 12037 | 10151 | 13238 | MOUSE | 5382 | 7977 | 8201 | 10152 | 389 | 347 | 2509 | 3245 |
| HUMAN | 9125 | 12037 | 10151 | 13238 | MYCTU | 563 | 1133 | 732 | 1456 | 189 | 46 | 617 | 852 |
| HUMAN | 9125 | 12037 | 10151 | 13238 | YEAST | 3191 | 4730 | 4256 | 4964 | 335 | 247 | 1267 | 1849 |
| MOUSE | 5382 | 7977 | 8201 | 10152 | ARATH | 5038 | 7352 | 7225 | 10002 | 253 | 166 | 1096 | 1515 |
| MOUSE | 5382 | 7977 | 8201 | 10152 | ECOLI | 2267 | 2228 | 2525 | 3288 | 205 | 56 | 688 | 949 |
| MOUSE | 5382 | 7977 | 8201 | 10152 | HUMAN | 9125 | 12037 | 10151 | 13238 | 389 | 347 | 2509 | 3245 |
| MOUSE | 5106 | 7571 | 7789 | 9644 | MOUSE | 276 | 406 | 412 | 508 | 189 | 199 | 1573 | 1961 |
| MOUSE | 5382 | 7977 | 8201 | 10152 | MYCTU | 563 | 1133 | 732 | 1456 | 146 | 38 | 527 | 711 |
| MOUSE | 5382 | 7977 | 8201 | 10152 | YEAST | 3191 | 4730 | 4256 | 4964 | 252 | 185 | 1065 | 1502 |
| MYCTU | 563 | 1133 | 732 | 1456 | ARATH | 5038 | 7352 | 7225 | 10002 | 119 | 38 | 407 | 564 |
| MYCTU | 563 | 1133 | 732 | 1456 | ECOLI | 2267 | 2228 | 2525 | 3288 | 94 | 30 | 281 | 405 |
| MYCTU | 563 | 1133 | 732 | 1456 | HUMAN | 9125 | 12037 | 10151 | 13238 | 189 | 46 | 617 | 852 |
| MYCTU | 563 | 1133 | 732 | 1456 | MOUSE | 5382 | 7977 | 8201 | 10152 | 146 | 38 | 527 | 711 |
| MYCTU | 536 | 1074 | 701 | 1383 | MYCTU | 27 | 59 | 31 | 73 | 21 | 12 | 89 | 122 |
| MYCTU | 563 | 1133 | 732 | 1456 | YEAST | 3191 | 4730 | 4256 | 4964 | 102 | 38 | 405 | 545 |
| YEAST | 3191 | 4730 | 4256 | 4964 | ARATH | 5038 | 7352 | 7225 | 10002 | 189 | 167 | 803 | 1159 |
| YEAST | 3191 | 4730 | 4256 | 4964 | ECOLI | 2267 | 2228 | 2525 | 3288 | 160 | 59 | 507 | 726 |
| YEAST | 3191 | 4730 | 4256 | 4964 | HUMAN | 9125 | 12037 | 10151 | 13238 | 335 | 247 | 1267 | 1849 |
| YEAST | 3191 | 4730 | 4256 | 4964 | MOUSE | 5382 | 7977 | 8201 | 10152 | 252 | 185 | 1065 | 1502 |
| YEAST | 3191 | 4730 | 4256 | 4964 | MYCTU | 563 | 1133 | 732 | 1456 | 102 | 38 | 405 | 545 |
| YEAST | 3022 | 4494 | 4038 | 4715 | YEAST | 169 | 236 | 218 | 249 | 109 | 147 | 557 | 813 |

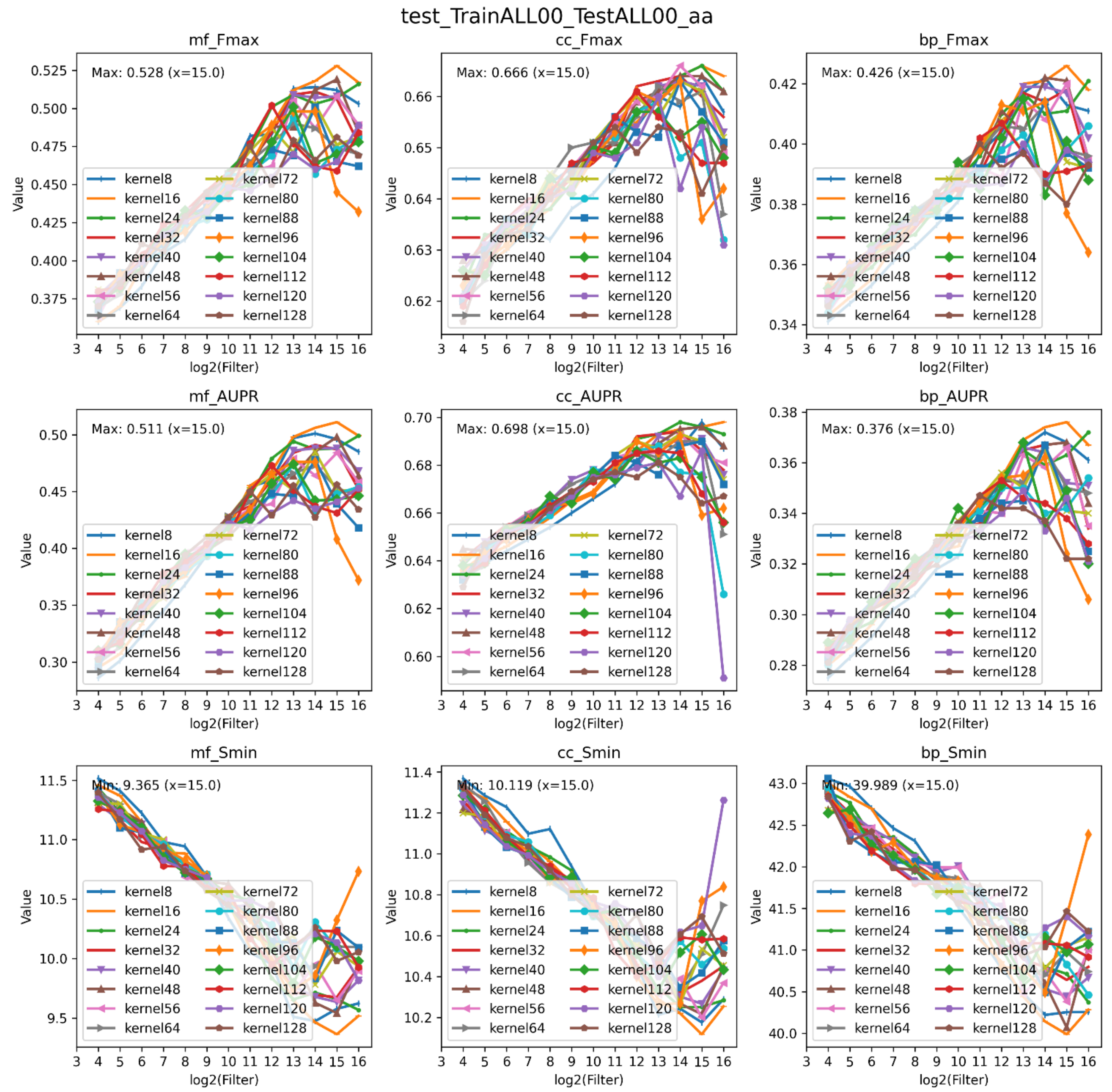

**Figure S1.** The evaluation results using the SwissProt dataset, utilizing primary sequences (aa) as input. It includes nine subfigures, each illustrating performance metrics across three subontologies: MFO, Cellular CCO, and BPO. The metrics evaluated are F*max*, AUPR, and S*min*. The horizontal axis of each subfigure represents the logarithmic scale of the filter size, while the vertical axis indicates the metric scores. Each plot within the figure correlates to the same kernel, providing a comprehensive overview of the model's performance across different parameters and ontologies.

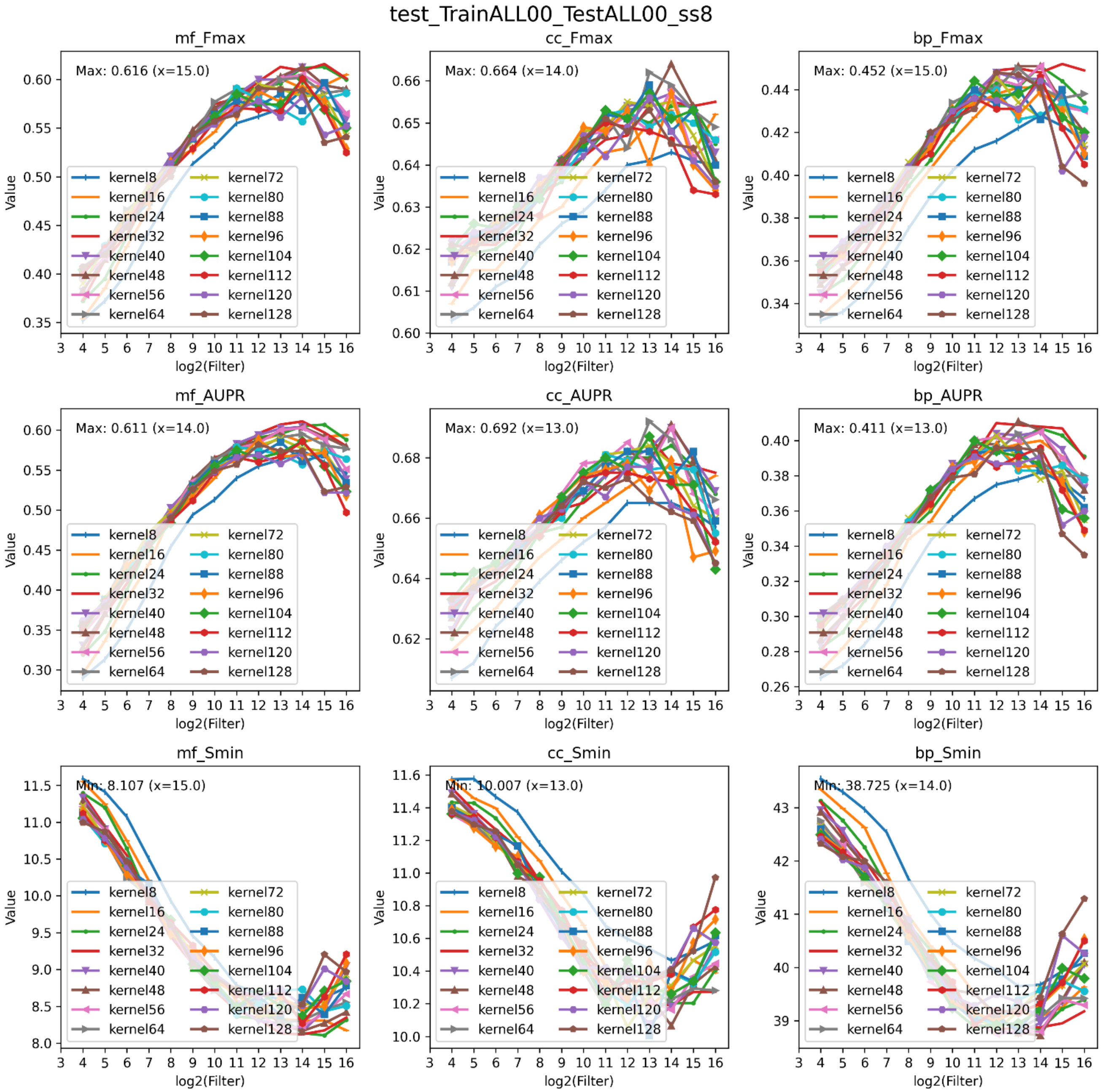

**Figure S2.** The evaluation results using the SwissProt dataset, utilizing secondary structure (ss8) as input. It includes nine subfigures, each illustrating performance metrics across three subontologies: MFO, Cellular CCO, and BPO. The metrics evaluated are F*max*, AUPR, and S*min*. The horizontal axis of each subfigure represents the logarithmic scale of the filter size, while the vertical axis indicates the metric scores. Each plot within the figure correlates to the same kernel, providing a comprehensive overview of the model's performance across different parameters and ontologies.

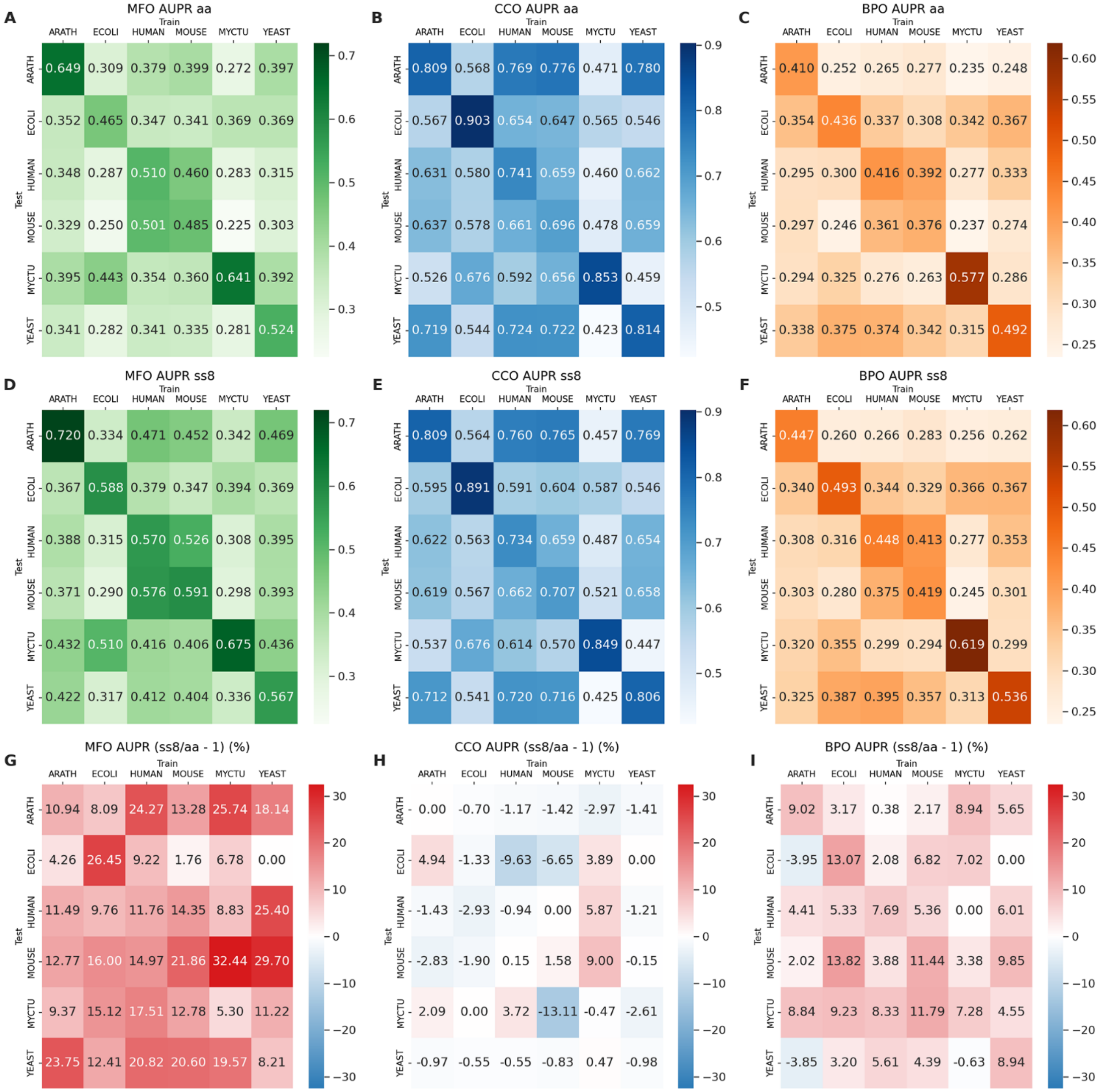

**Figure S3.** The performance of AUPR scores in predicting GO functional annotations across species is depicted through heatmaps, utilizing models that have been trained and tested among six different species: Arabidopsis thaliana (ARATH), Escherichia coli (ECOLI), Homo sapiens (HUMAN), Mus musculus (MOUSE), Mycobacterium tuberculosis (MYCTU), and Saccharomyces cerevisiae (YEAST). (A) and (D) present MFO results based on model-aa and model-ss8 respectively; (B) and (E) show CCO results; and (C) and (F) illustrate BPO outcomes. Darker shades in the color gradients indicate higher metrics scores, reflecting greater prediction accuracy. Each matrix cell provides a metrics score for a model trained on the species denoted at the top and tested on the species labeled on the side. (G) depicts the percentage increased performance from model-aa to model-ss8 in MFO, similarly, (H) and (I) represent the increments in CCO and BPO, respectively. Red indicates the percentage of increase, while blue represents the percentage of decrease.

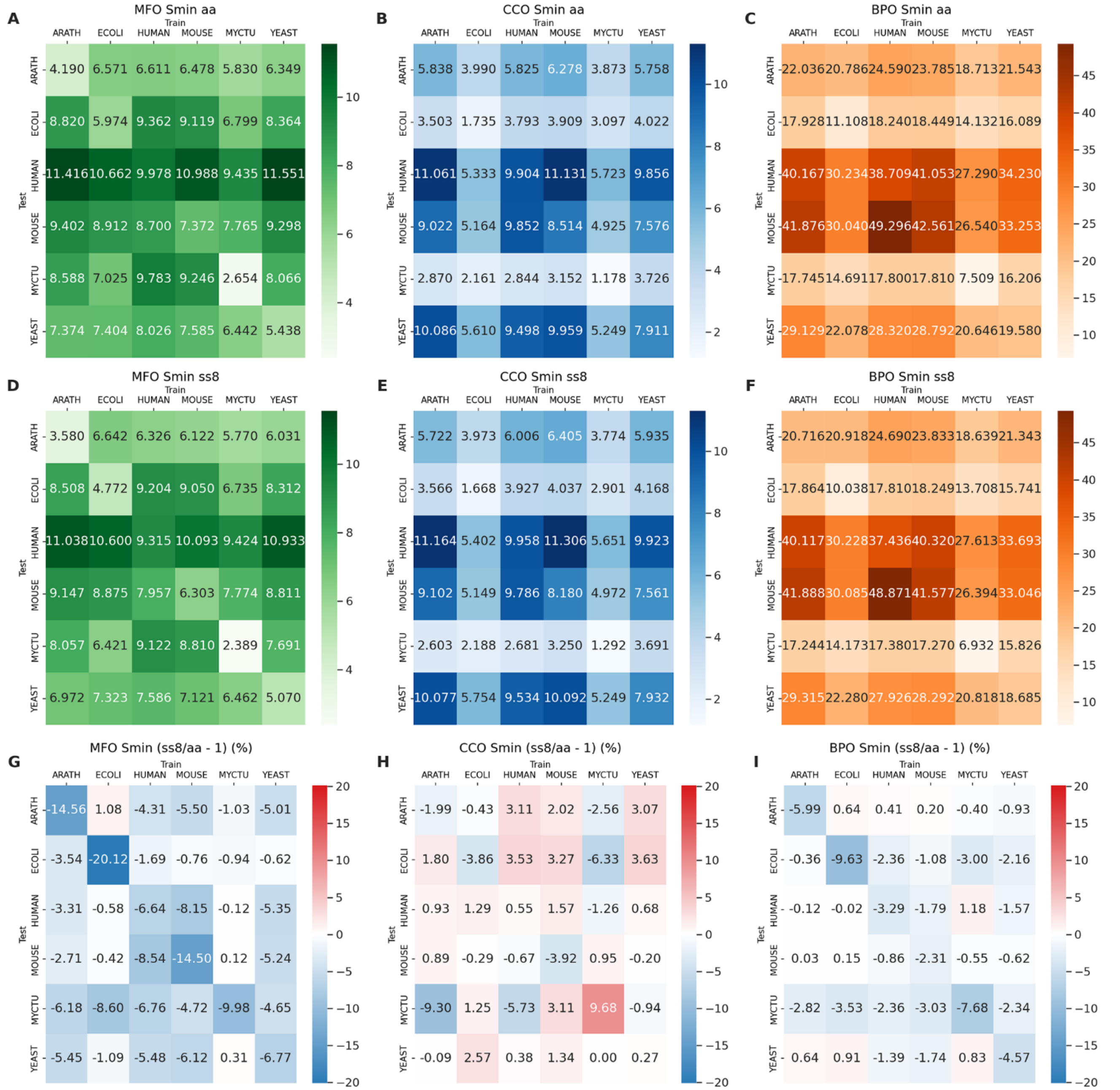

**Figure S4.** The performance of S*min* scores in predicting GO functional annotations across species is depicted through heatmaps, utilizing models that have been trained and tested among six different species: Arabidopsis thaliana (ARATH), Escherichia coli (ECOLI), Homo sapiens (HUMAN), Mus musculus (MOUSE), Mycobacterium tuberculosis (MYCTU), and Saccharomyces cerevisiae (YEAST). (A) and (D) present MFO results based on model-aa and model-ss8 respectively; (B) and (E) show CCO results; and (C) and (F) illustrate BPO outcomes. Darker shades in the color gradients indicate higher metrics scores, reflecting greater prediction accuracy. Each matrix cell provides a metrics score for a model trained on the species denoted at the top and tested on the species labeled on the side. (G) depicts the percentage increased performance from model-aa to model-ss8 in MFO, similarly, (H) and (I) represent the increments in CCO and BPO, respectively. Red indicates the percentage of increase, while blue represents the percentage of decrease.

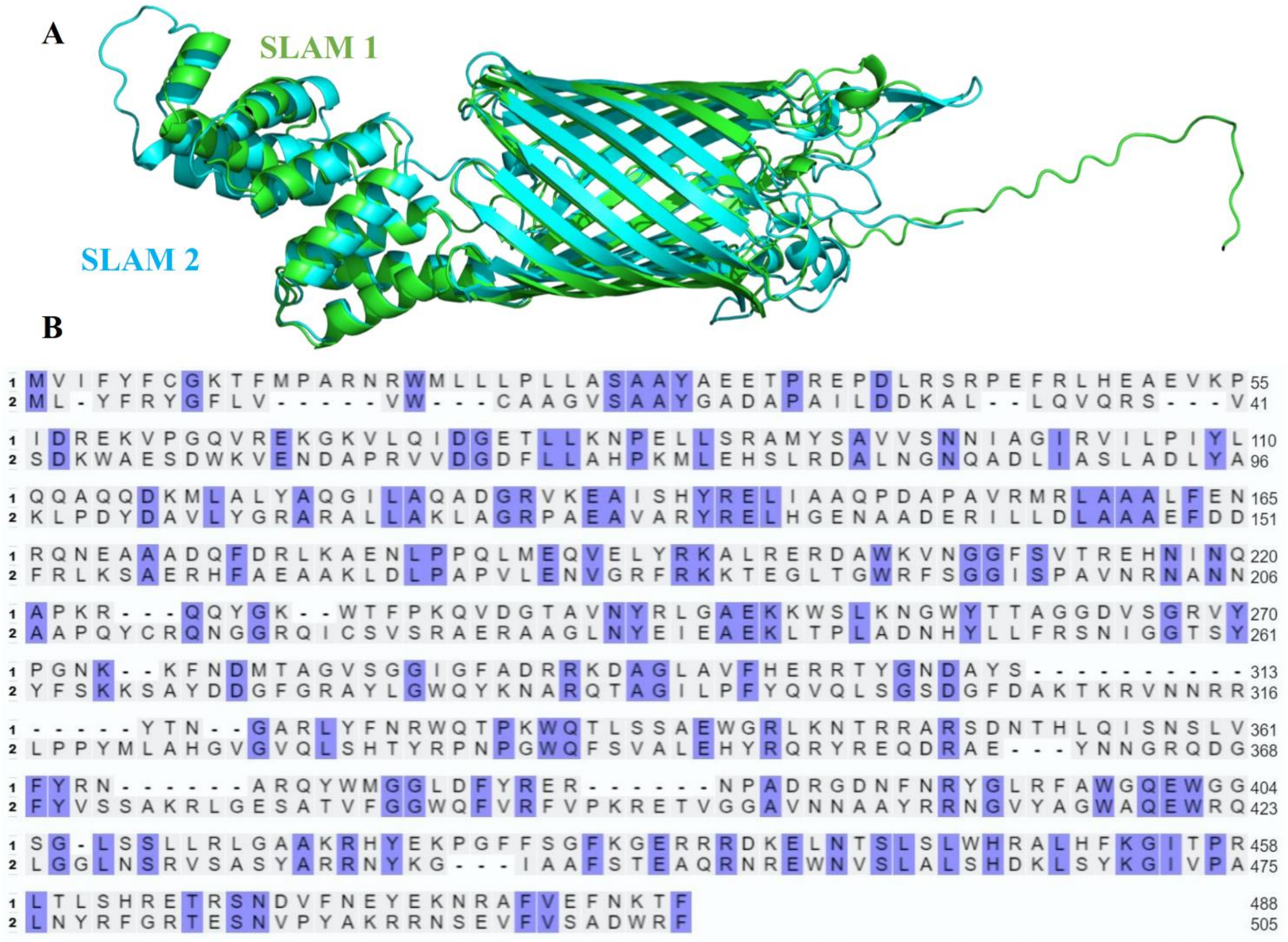

**Figure S5.** (A) The schematic diagram of the AlphaFold2 3D spatial structure prediction for SLAM1 and SLAM2 proteins (Surface Lipoprotein Assembly Modifier). Both comprise a Beta barrel and several Alpha helices, with highly similar secondary and tertiary structures. (B) The sequence alignment results for SLAM1 and SLAM2, with highlighted letters indicating identical amino acids. The sequence difference between the two is very large, with only about 25% sequence identity.
